# Induction of common bean *OVATE Family Protein 7* (*PvOFP7*) promotes resistance to common bacterial blight

**DOI:** 10.1101/2024.01.26.577399

**Authors:** Charlotte Gaudin, Anne Preveaux, Nathan Aubineau, Damien Le Goff, Marie-Agnès Jacques, Nicolas W.G. Chen

**Affiliations:** Univ Angers, Institut Agro, INRAE, IRHS, SFR QUASAV, F-49000 Angers, France

**Author notes:** Author for correspondence: *Nicolas W.G. Chen, Email:.

**Keywords:** Common bacterial blight, *Phaseolus vulgaris*, disease, resistance, *Xanthomonas*, Heat shock proteins, cell wall

## Abstract

Common bacterial blight of bean (CBB) is a devastating seed-transmitted disease caused by *Xanthomonas phaseoli* pv. *phaseoli* and *Xanthomonas citri* pv. *fuscans* on common bean (*Phaseolus vulgaris* L.). The genes responsible for CBB resistance are largely unknown. moreover, the lack of reproducible and universal transformation protocol limits the study and improvement of genetic traits in common bean. We produced *X. phaseoli* pv. *phaseoli* strains expressing artificially-designed Transcription-Activator Like Effectors (dTALEs) to target 14 candidate genes and performed *in planta* assays in a susceptible common bean genotype to analyse if the transcriptional induction of these genes could confer resistance to CBB. Induction of *PvOFP7*, *PvAP2-ERF71* and *PvExpansinA17* resulted in CBB symptom reduction. In particular, *PvOFP7* induction led to strong symptom reduction, linked to reduced bacterial growth *in planta* at early colonisation stages. RNA-Seq analysis revealed up-regulation of cell wall formation and primary metabolism, and major down-regulation of Heat Shock Proteins. Our results demonstrate that PvOFP7 is contributes to CBB resistance, and underline the usefulness of dTALEs for highlighting genes of quantitative activity.

## INTRODUCTION

Plants possess an immune system to defend themselves against a large array of pathogens (Jones and Dangl, 2006). The first layer of this immune system corresponds either to the extracellular perception of microbial- or pathogen-associated molecular patterns (MAMPs or PAMPs) through pattern-recognition receptors (PRRs) present at the surface of the plant cell, or to the intercellular perception of microbial effectors through proteins belonging to the nucleotide-binding site leucine-rich repeat (NLR) family (Dodds and Rathjen, 2010; Ngou *et al*., 2022). Following perception, both PAMP-triggered immunity (PTI) and effector-triggered immunity (ETI) initiate complex signalling pathways usually involving a burst of reactive oxygen species (ROS), calcium influx and mitogen-activated protein kinase (MAPK) cascades, leading to the activation of multiple transcription factors (TF) and to the expression of defense-related genes (Peng *et al*., 2018; Tsuda and Katagiri, 2010; Yuan *et al*., 2021). These pathways are associated to the biosynthesis of defense-related phytohormones such as salicylic acid (SA), jasmonic acid (JA) or ethylene (ET) and ultimately lead to mechanisms aiming at limiting pathogen propagation (Pieterse *et al*., 2012; Checker *et al*., 2018). These mechanisms may include strengthening of the plant cell wall through callose deposition and/or production of antimicrobial compounds. The ETI response is often more intense than PTI and usually results in a hypersensitive response (HR) associated to localized cell death to halt pathogen progression (Balint-Kurti, 2019; Yuan *et al*., 2021).

Legumes constitute a major food group presenting both high nutritional properties and low negative environmental impact (Willett *et al*., 2019; Foyer *et al*., 2016). As such, legumes contribute both to healthy people and a healthy planet (Uebersax *et al*., 2022). Among legumes, common bean (*Phaseolus vulgaris* L.) is an important crop for direct human consumption that is used worldwide for its edible pods and seeds (Bitocchi *et al*., 2017). In addition to having recognized human health benefits and nutritional composition (Didinger and Thompson, 2022), common bean is considered an affordable species for sustainable agriculture (Uebersax *et al*., 2022).

Common bacterial blight of bean (CBB) is a devastating seed-transmitted disease caused by two phylogenetically distant bacterial species, *Xanthomonas phaseoli* pv. *phaseoli* and *Xanthomonas citri* pv. *fuscans* that induce identical symptoms on common bean (Chen *et al*., 2021). This disease represents an important economic and agricultural threat to both edible bean production and bean seed industry worldwide. Therefore, it is essential to better understand the determinants of resistance to CBB in order to achieve durable common bean production. Resistance to CBB has been extensively studied (Singh and Miklas, 2015; Singh and Schwartz, 2010) and appears to be a complex process involving numerous quantitative trait loci (QTLs), but the genes responsible for this trait remain largely unknown.

More generally, a limiting aspect to the study and improvement of genetic traits in common bean is the lack of reproducible and universal transformation protocol. Indeed, although successful transformation has been achieved in few cases (Bonfim *et al*., 2007; Aragão *et al*., 1998; Collado *et al*., 2015), protocols remain largely genotype-dependent and both transformation and regeneration rates are still extremely low (Veltcheva *et al*., 2005; Hnatuszko-Konka *et al*., 2019; Hnatuszko-konka *et al*., 2014). Thus, except for hairy-root systems (Voß *et al*., 2022; Estrada-Navarrete *et al*., 2007), gene editing methods such as CRISPR/Cas9 have yet to be democratized for common bean (Toili, 2022). Transient expression via *Agrobacterium* (Richard et al., 2021) and virus-induced gene silencing (VIGS) methods have been developed using the bean pod mottle virus (Pflieger *et al*., 2014; Zhang *et al*., 2013). However, for VIGS to work, the tested genotypes must be susceptible to the virus, and the target gene expression or silencing takes days, which is often incompatible with disease phenotyping assays.

Bacteria from the *Xanthomonas* genus possess the ability of directly manipulating their host transcriptome by using type III effectors named Transcription-Activator Like Effectors (TALEs) able to bind to specific DNA sequences in the promoter region of target genes and recruit the host cell’s transcriptional machinery to induce gene expression (Boch and Bonas, 2010; Perez-Quintero and Szurek, 2019). The central part of the TALE protein is composed of a variable number of repeats of 34 amino acids each, resulting in a helical structure that allows the TALE to wrap around the DNA double helix (Mak *et al*., 2013; Deng *et al*., 2012; Mak *et al*., 2012). Within each repeat, the doublet of amino acids at positions 12 and 13, called repeat variable diresidue (RVD), determines the binding specificity to a given nucleotide (Moscou and Bogdanove, 2009; Boch *et al*., 2009).

The TALE code linking RVDs to target nucleotides has led to the development of a cloning kit enabling *in vitro* construction of designer TALEs (dTALEs) of any chosen RVD order, which can be used to induce any desired gene within an eukaryotic genome (Geißler *et al*., 2011). Here, we used dTALEs as tools to study 14 candidate genes presenting different expression patterns during CBB infection in resistant versus susceptible common bean genotypes (Foucher *et al*., 2020). We produced *X. phaseoli* pv. *phaseoli* strains expressing dTALEs targeting the promoter region of each candidate gene and performed *in planta* assays to analyse if the transcriptional induction of these genes in a susceptible common bean genotype could confer resistance to CBB.

## MATERIALS AND METHODS

### Bacterial strains and growing conditions

Bacterial strains used in this study are wild-type and dTALE-complemented versions of *Xanthomonas phaseoli* pv. *phaseoli* strain 6546R (Briand *et al*., 2021), a spontaneous rifamycin-resistant derivative of strain CFBP 6546. All strains were grown for 48h at 28°C on Trypticase Soy Agar (TSA) medium (17.0 g.l^-1^ pancreatic digest of casein, 3.0 g.l^-1^ enzymatic digest of soya bean, 5.0 g.l^-1^ NaCl, 2.5 g.l^-1^ K_2_HPO_4_, 2.5 g.l^-1^ glucose, 15 g.l^-1^ agar, pH 7.3 at 25°C), then for 24h at 28°C on a 1/10 dilution of TSA medium (except for agar kept at 15 g.l^-1^) to obtain fresh bacterial cultures for pathogenicity assays. When necessary, antibiotics were added to the medium at the following final concentrations: rifamycin, 50 µg.ml^-1^ and gentamycin, 20 µg.ml^-1^.

### Construction of dTALEs

Candidate genes potentially involved in common bean resistance to CBB were retrieved from Foucher *et al*. (2020). The promoter sequence of each candidate gene was analysed with TAL Effector Targeter, a bioinformatic tool that can predict optimal dTALE binding sites (Doyle *et al*., 2012; Cermak *et al*., 2011). The specificity of target sequences was verified with BLASTn (Evalue < 1) on both the genome of common bean genotype JaloEEP558 (Altschul *et al*., 1990; Foucher *et al*., 2020) and the common bean reference genome (v2.1, DOE-JGI and USDA-NIFA, http://phytozome.jgi.doe.gov/; Schmutz et al. 2014). The dTALEs were built using the Golden TAL technology (Geißler *et al*., 2011) and expressed in the Golden Gate-compatible vector pSKX1 (Streubel *et al*., 2013). The dTALE constructions, or the empty vector (EV) used as control, were introduced into strain 6546R by electroporation. Rifamycin- and gentamycin-resistant clones were selected upon plating on 1/10 TSA medium supplemented with appropriate antibiotics. For each dTALE, *in vitro* growth assays were performed on three different clones, in order to choose the clone that had the closest growth profile compared to the strain carrying an empty vector.

### Pathogenicity assays

Plants from the CBB-susceptible common bean (*Phaseolus vulgaris* L.) genotypes JaloEEP558, Contender, Facila, Flavert and Linex were cultivated in a growth chamber at 23/20°C (day/night) with 80% relative humidity and a photoperiod of 16 h. Plants were watered every two days, with water for the first 10 days, then with a nutrient solution of N-P-K (15–10-30) until the end of the assay. One day before inoculation, temperature and humidity were increased to 28/25°C and 95% relative humidity to optimize infection. Inoculation was performed by rubbing the first leaves of 8-day-old plants using bacterial suspensions at 1 × 10^8^ cfu.ml^-1^ for symptom quantification, or 1 × 10^6^ cfu.ml^-1^ for bacterial population measurements, as described in Foucher *et al*. (2021), except that inoculates were rubbed using a dauber bottle (Vaessen creative, ref: SKU:7005-009) instead of gloved finger. Water-inoculated plants were used as controls. Two days after inoculation, relative humidity was decreased back to 80%, but temperature was maintained at 28/25°C. *In planta* bacterial populations were evaluated at 0, 4, 7, and 11 days post inoculation (dpi). Each sample corresponded to twelve 1.2cm-diameter disks collected from the two first leaves of each plant (6 disks per leaf). Samples were put in leak-proof plastic bags, ground with a rolling pin and resuspended in 2ml of sterile water. Suspensions of appropriate dilutions were plated on 1/10 TSA medium supplemented with appropriate antibiotics, then kept at 28°C for 72h before counting. Symptoms were quantified 10 dpi by machine learning-based imaging using Ilastik v.1.3.3 (Berg *et al*., 2019) as previously described (Foucher *et al*., 2021).

### RNA isolation and RT-qPCR assays

To obtain homogeneously inoculated plant tissues, leaflets from the first trifoliate leaves (stage V1) were detached from the plant and vacuum-infiltrated for 1 min into bacterial suspensions at 1 × 10^8^ cfu.ml^−1^ or pure sterile distilled water as control. Infiltrated leaflets were maintained in Petri dishes by dipping the petiole in water agar (0.7%) and incubated at 28°C/25°C (day/ night) with a photoperiod of 16 h. Sampling was performed 48 hours after inoculation. Each sample corresponded to six 1.2 cm-diameter leaf disks collected from three leaflets (two disks per leaflet) and immediately frozen in liquid nitrogen, then conserved at −80°C. Samples were ground in fine powder using a ball mill for 2×30s at 30Hz. Total RNA was isolated using the NucleoSpin RNA Plus Mini kit (Macherey-Nagel, Hoerdt, France) following manufacturer’s recommendations.

Reverse transcription, qPCR primers design and qPCR assays were performed as described in Foucher *et al*. (2020). Relative expression levels were calculated using the 2^(–ΔΔCt)^ method (Vandesompele *et al*., 2002), and normalized to reference genes *EF1-α* (Phvul.004G060000), *Act11* (Phvul.008G011000) and *IDE* (Phvul.001G133200). Each experiment was performed using three biological samples as replicates, with two technical replicates per sample.

### RNA-Seq experiments

RNA extraction was performed following the same procedure as described for RT-qPCR assays, using three independent biological samples per modality. RNA-Seq was performed at the Beijing Genomics Institute (BGI), Hong Kong, China. RNA-Seq libraries were prepared using the DNBSEQ Transcriptome protocol. Quality control was done using Agilent 2100 chips. Paired-end RNA sequencing (2 x 100 bp) was performed on a DNBSEQ-G400 sequencer. After filtering and quality control using Phred+33, this led to around 24 million clean paired-end reads per sample, with Q20% above 98%. Total reads were mapped to the common bean reference genome (Schmutz *et al*., 2014) and counted using Salmon v1.5.1 (Patro *et al*., 2017).

### Analysis of differentially-expressed genes

Analysis of RNA-Seq data was performed using the R pipeline AskoR v0.1 (Carvalho *et al*., 2021) on the GeneOuest bioinformatics Galaxy platform (Galaxy GenOuest, https://www.genouest.org). AskoR performs a suite of statistical analyses including filtering and normalization of raw counts into count per million (CPM) mapped reads, Principal Component Analysis (PCA), differential gene expression analysis using EdgeR v3.32.1 with default parameters (Robinson *et al*., 2009), as well as different graphical outputs such as volcano plots and heatmaps (Supplemental figure 1). Differentially-expressed genes (DEG) were retrieved using the EV strain as control, with false discovery rate (FDR) threshold of 0.05 and |Log_2_(FC)| > 1. Lists of common bean genes corresponding to transcription factors, transcription regulators and kinases (including receptor-like kinases) were retrieved from the iTAK database (http://itak.feilab.net) (Zheng *et al*., 2016). The complete set of genes encoding nucleotide-binding leucine-rich repeat (NL) proteins was retrieved according to Richard et al. (2017). Enrichment of functional categories were performed using the Wilcoxon rank sum test and PageMan functions available in MapMan v3.6.0RC1 (Usadel *et al*., 2005; Thimm *et al*., 2004), which was also used for visualisation of DEGs in different functional pathways.

### Phylogenetic analysis

Amino acid sequences of 19 OVATE Family Protein (OFP) genes from *Arabidopsis thaliana* (Liu *et al*., 2014) and 20 from common bean (Schmutz et al. 2014) were retrieved and aligned using MAFFT v.1.3.6 in Geneious v.9.1.8 (https://www.geneious.com). Phylogenetic analysis was made on the conserved OVATE domain (Liu *et al*., 2014). A maximum likelihood phylogenetic tree was generated using PhyML v.2.2.3 in Geneious v.9.1.8 (https://www.geneious.com) and 100 bootstrap iterations. The tree was visualised and edited with MEGA v.3.6.0 (Kumar *et al*., 2008).

## RESULTS

### Candidate gene screening identified three genes linked to CBB symptom reduction

To search for genes involved in common bean resistance to CBB, we used *X. phaseoli* pv. *phaseoli* 6546R strains complemented with dTALEs to induce the expression of 14 candidate genes in the susceptible bean genotype JaloEEP558 (Supplemental table 1). These genes were chosen based on their significantly higher expression in resistant *vs* susceptible bean genotypes 48h after inoculation (Foucher *et al*., 2020). Of the 14 dTALE-complemented strains, 13 led to successful induction of corresponding target gene (Supplemental figure 2). Among those, pathogenicity assays highlighted three genes (Phvul.004G157600, Phvul.007G207600 and Phvul.009G057100) whose induction was related to significant symptom reduction compared to the EV strain (Fig. 1; Supplemental figure 3). These genes were respectively induced by dTALEs ERF_A, ExpansinA17_A and OFP7_A (Supplemental table 1). Phvul.004G157600 encoded the APETALA2/ETHYLENE RESPONSIVE FACTOR (AP2/ERF) protein PvAP2-ERF71 (Kavas *et al*., 2015), while Phvul.007G207600 and Phvul.009G057100 encoded the expansin PvExpansinA17 and the OVATE family protein (OFP) PvOFP7, respectively, based on homology with *Arabidopsis*.

**Fig. 1.**
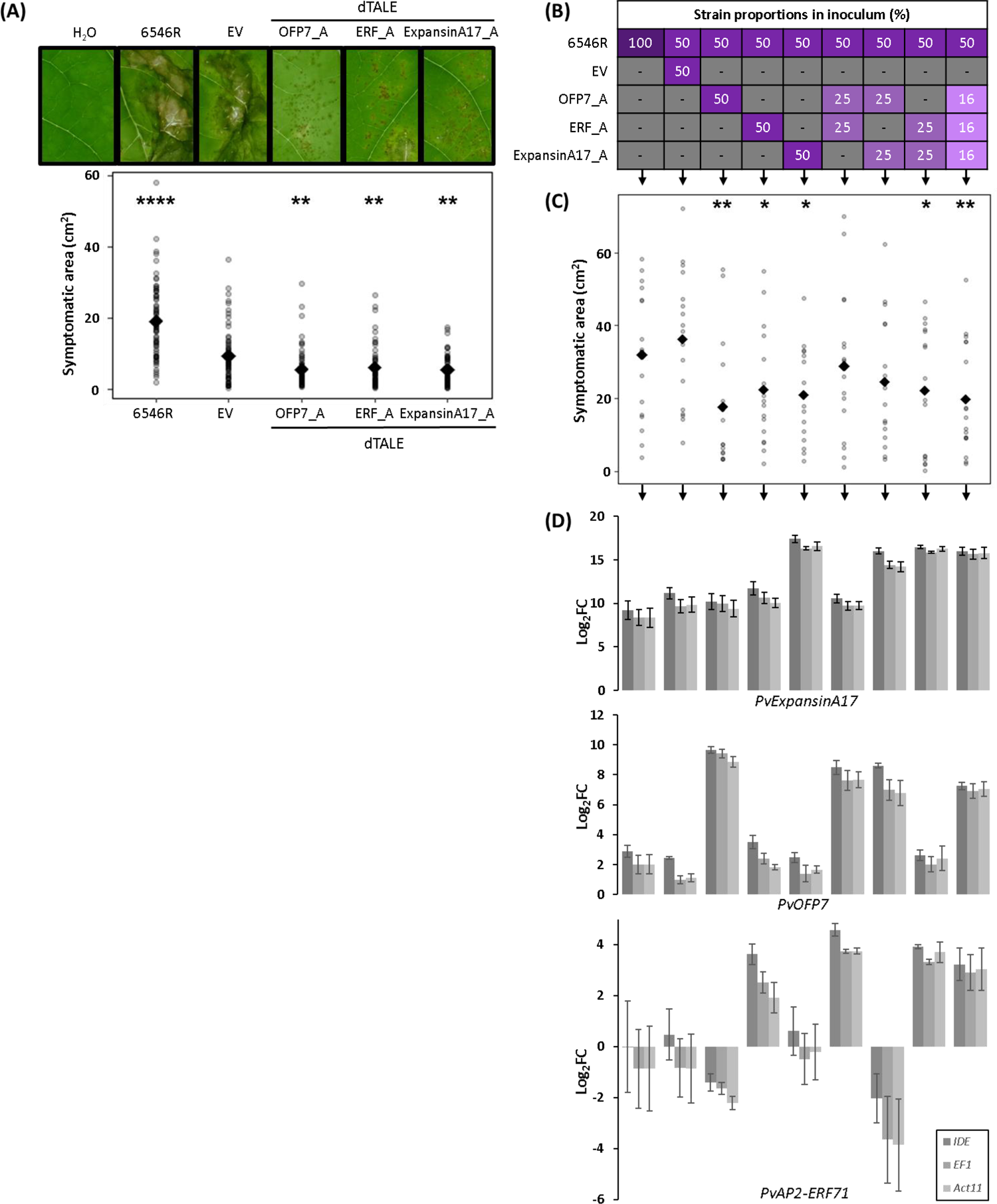
CBB symptom reduction observed after induction of three candidate genes. First leaves were inoculated by strain 6546R, or strains expressing dTALEs targeting common bean genes *PvOFP7* (strair OFP7_A), *PvERF* (strain ERF A), *PvExpansinAl’/* (strain ExpansinA17_A), or carrying an empty vectoi (EV). **(A)** Average symptomatic area quantified 10 days after inoculation of individual strains. Results presentee here combined four independent experiments (n = 6‘ plants per treatment). (B) Table presenting bacterial mixtures used for experiments described in (C) and (D) Mixtures were made up of 50% of strain 6546R and 50°/< of dTALE-bearing strains, each at equal concentrations **(C)** Average symptomatic area quantified 10 days aftei co-inoculation of strain mixtures described in table (B) Results cumulated data from two independent experiments (n = 17 plants per treatment). **(D)** Expression levels of *PvExpansinAl7, PvOFP7* anc *PvAP2-ERF71* measured by qRT-PCR at 2 days aftei vacuum infiltration of strain mixtures presented in table (B). Log_2_(FC) were calculated using the 2^( AACT)^ methoc (n = 3 samples per treatment). For all experiments, inoculum concentration was 1 × 10^2^ cfu.ml” f For (A) and (C), each dot represents one sample, while squares correspond to the mean for eacf treatment. Kruskal Wallis test indicated significant effeci (PO.05) for both (A) and (C). The number of stars refers to significantly different values compared to the EV (A^ or the 1:1 mixture of strains 6546R and EV (seconc column) in (C) after Dunn’s test (*, P< 0.05; **, P< 0.01 ***, P< 0.001; **** P< 0.0001).

For these three genes, significant symptom reduction was still observed for dTALE-carrying strains co-inoculated at equal concentrations with strain 6546R, compared to the 6546R + EV co-inoculation (Fig. 1B). This strongly suggested that the observed reduction of symptoms was indeed due to effective resistance. Intriguingly, symptom reduction was lost when co-inoculating strain 6546R with strains bearing OFP7_A together with either ExpansinA17_A or ERF_A, but was restored when co-inoculating the four strains together (Fig. 1B). This suggested negative regulatory relationships between the three genes. In accordance with this, co-induction of *PvOFP7* and *PvExpansinA17* resulted in down-regulation of *PvAP2-ERF71* (Fig. 1C). Together, these results unveiled potential regulators of common bean resistance to CBB, acting through complex and non-complementary regulatory interactions.

### Induction of *PvOFP7* promotes partial resistance to CBB

In the rest of this paper, we focused on *PvOFP7* because it was previously highlighted as being the most differentially-expressed gene between resistant and susceptible plants (Foucher *et al*., 2020), and was located in a QTL of resistance to CBB (Nodari et al. 1993), which was not the case for the other two candidates. PvOFP7 belonged to the OFP family of transcription factors, comprising 20 and 19 members in *P. vulgaris* and *Arabidopsis thaliana*, respectively (Supplemental figure 4). To validate that the observed decrease of CBB symptoms was indeed due to the induction of *PvOFP7* and not of an off-target in JaloEEP558 genome, we constructed two additional dTALEs (OFP7_B and OFP7_C) targeting different regions of the *PvOFP7* promoter (Fig. 2). For each dTALE, induction of *PvOFP7* was linked to more than 60% symptoms reduction compared to the EV strain (Fig. 3A-C). In addition, *in planta* bacterial populations were significantly reduced at 4 dpi (Fig. 3D). This observed *PvOFP7*-mediated resistance appeared partial and transitory as bacterial populations caught up with strain 6546R at 7 dpi.

**Fig. 2.**
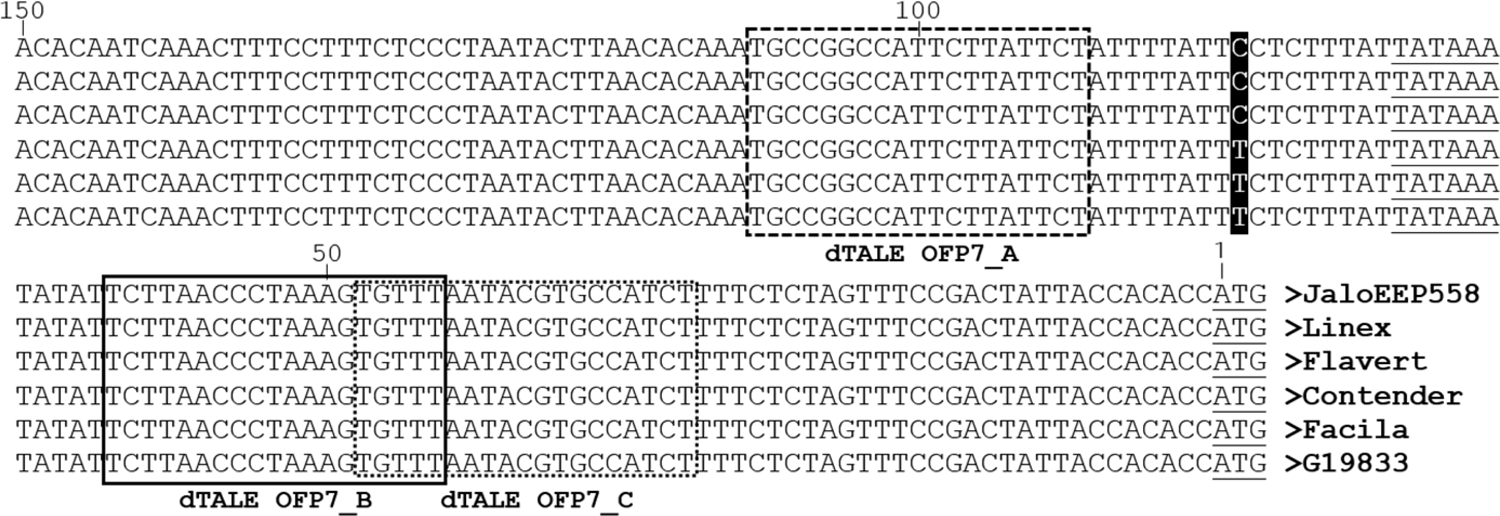
Alignment of *PvOFP7* promoter region in different common bean genotypes. The TATA box and start codon are underlined. Target sequences are framed using dashed lines for dTALE-OPF7_A, solid lines for dTALE-OPF7_B and dotted lines for dTALE-OPF7_C. One polymorphic nucleotide is indicated in black while other nucleotides are identical across all sequences.

**Fig. 3.**
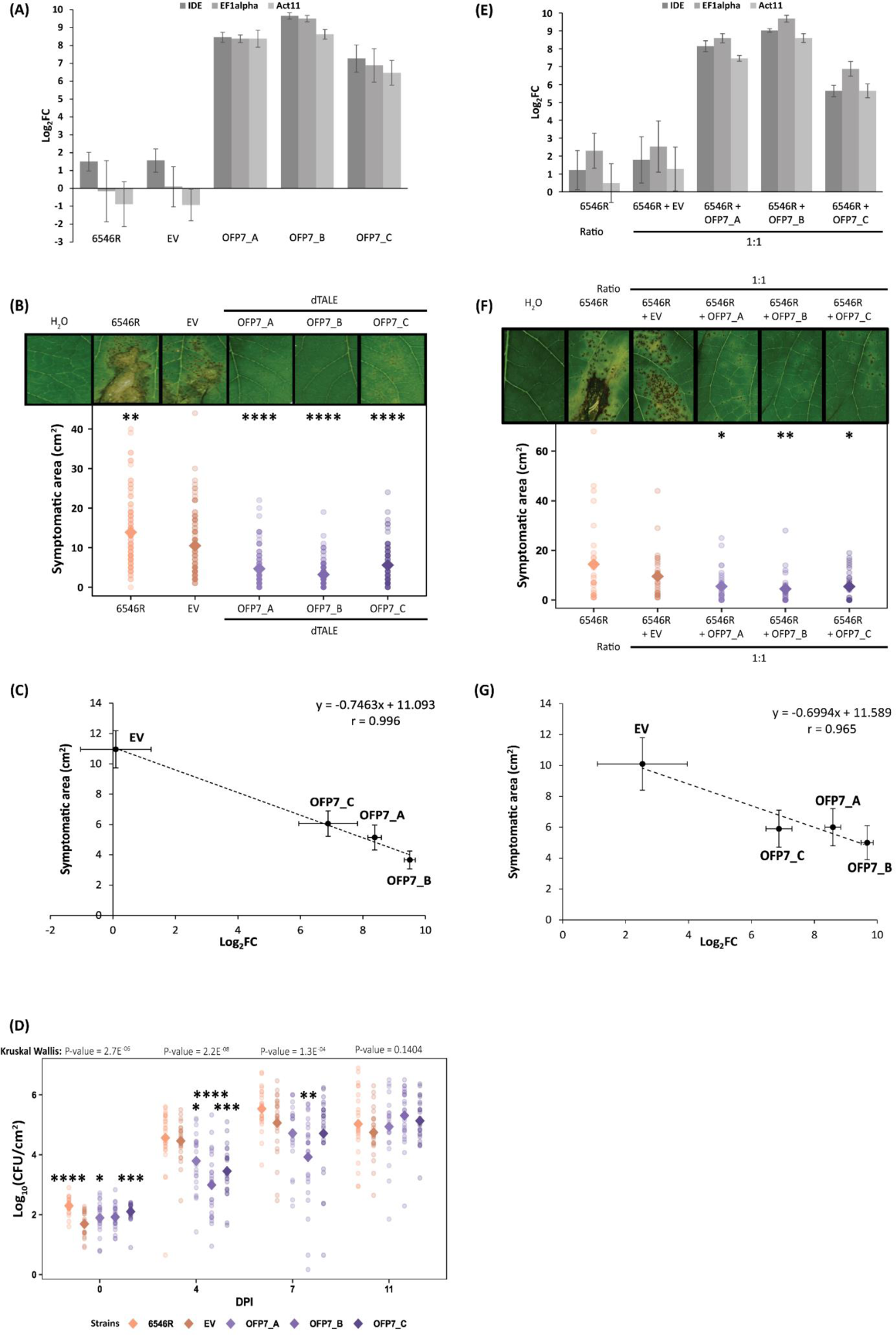
Induction of *PvOFP7* correlates with decreased susceptibility to common bacterial blight. **(A)** and **(E)** *PvOFP7* expression levels measured by qRT-PCR two days after vacuum-infiltration. Log_2_(FC) were calculated using the 2f^AACT)^ method. These results come from two independent experiments, each comprising three samples per treatment (n^_^6). (B) and (F) Average symptomatic area 10 days after inoculation of individual strains (n=100 plants per treatment) or mixtures of strains co-inoculated at a 1:1 ratio (n=30 plants per treatment), respectively. Each dot corresponds to one plant, while coloured squares correspond to the mean for each treatment. **(C)** and **(G)** Linear regressions between *in planta PvOFP7* expression levels and average symptomatic areas. (C) was done using the data presented in (A) and (B) while (G) used the data from (E) and (F). In both experiments, correlation coefficient r was significant (P<0.01) for n=4 observations. **(D)** Bacterial population sizes over time. Dunn’s test was performed separately at each sampling time. These results come from three independent experiments with 24 to 25 plants per condition. For (B), (G) and (D), Kruskal Wallis test indicated significant effect (p<0.05) of the treatments. The number of stars refers to significantly different values compared to the EV after Dunn’s test (*, P< 0.05; **, P< 0.01; ***, P< 0.001; ****, p< 0.0001).

Overall, the three dTALEs led to different levels of *PvOFP7* induction, symptoms and bacterial populations, with OFP7_B being the dTALE leading to the most significant differences compared to the EV. Remarkably, we observed a strong negative correlation (r = 0.996) between symptoms and *PvOFP7* induction levels (Fig. 3C). Such correlation was still observed for dTALE-carrying strains co-inoculated at a 1:1 ratio with strain 6546R (Fig. 3E-G), again indicating that *PvOFP7* induction was able to attenuate symptoms conferred by a wild-type strain. Altogether, these results indicated that the overexpression of *PvOFP7* contributed to dose-dependent reduction of CBB symptoms by impacting bacterial populations at the early stage of colonisation.

Moreover, the promoter region of *PvOFP7* was highly conserved across different common bean genotypes (Fig. 2). Using OFP7_B, *PvOFP7* induction led to decreased symptoms in four more or less susceptible genotypes, although not significant for Facila (Fig. 4; Supplemental figure 5) suggesting that *PvOFP7*-mediated resistance to CBB was ubiquitous in common bean.

**Fig. 4.**
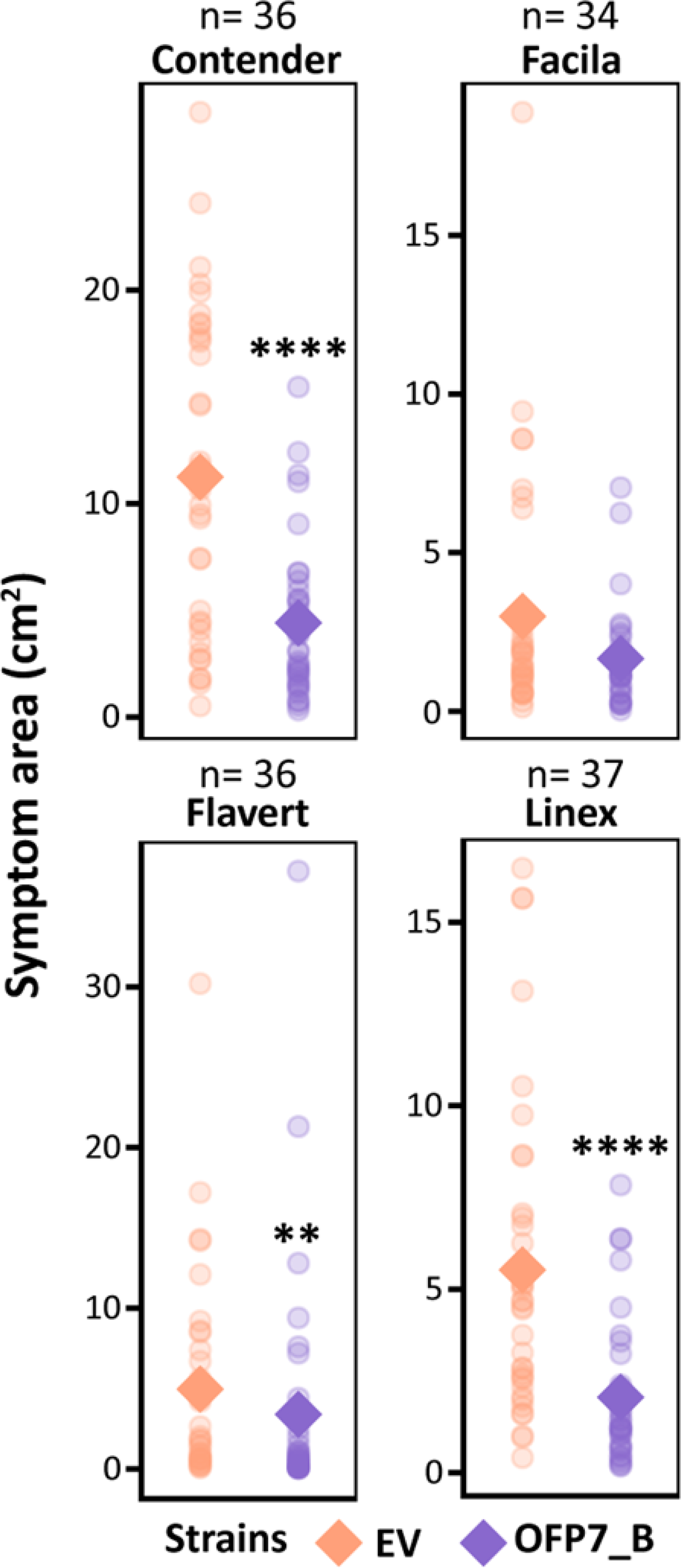
*PvOFP7* induction confers partial resistance to CBB in four additional common bean accessions. Results presented here correspond to symptomatic area at 10 dpi assessed in two independent experiments, with a total of 34 to 37 plants per condition. Kruskal Wallis test indicated significant effect (P<0.05) of the treatments, except for Facila. The number of stars refers to significantly different values compared to the EV after Dunn’s test (*, P< 0.05; **, P< 0.01; ***, P< 0.001; ****, p< 0.0001).

### Impact of *PvOFP7* induction on the common bean transcriptome

To study the metabolic pathways associated to *PvOFP7* induction in common bean, we performed an RNA-Seq analysis 48h after leaf inoculation of JaloEEP558. Principal Component Analysis showed that plants inoculated with the EV or 6546R strains formed a distinct group compared to plants inoculated with the strains carrying the dTALEs (Supplemental figure 6). In accordance with this, only four DEGs were found in plants inoculated with strain 6546R compared to EV strains indicating that the plasmid used for dTALE expression had almost no impact on the plant transcriptome (Supplemental table 2). For the strains carrying dTALEs, 2008 to 5648 DEGs were found (Supplemental figure 7), among which 952 were common to the three dTALEs (Fig. 5). Therefore, we considered that these 952 DEG were related to pathways impacted by the transcriptional induction of *PvOFP7*. Among these 952 DEG, 549 were down-regulated while 403 were up-regulated. This included *PvOFP7*, for which RNA-Seq expression levels correlated well with qRT-PCR results, (Supplemental figure 8).

**Fig. 5.**
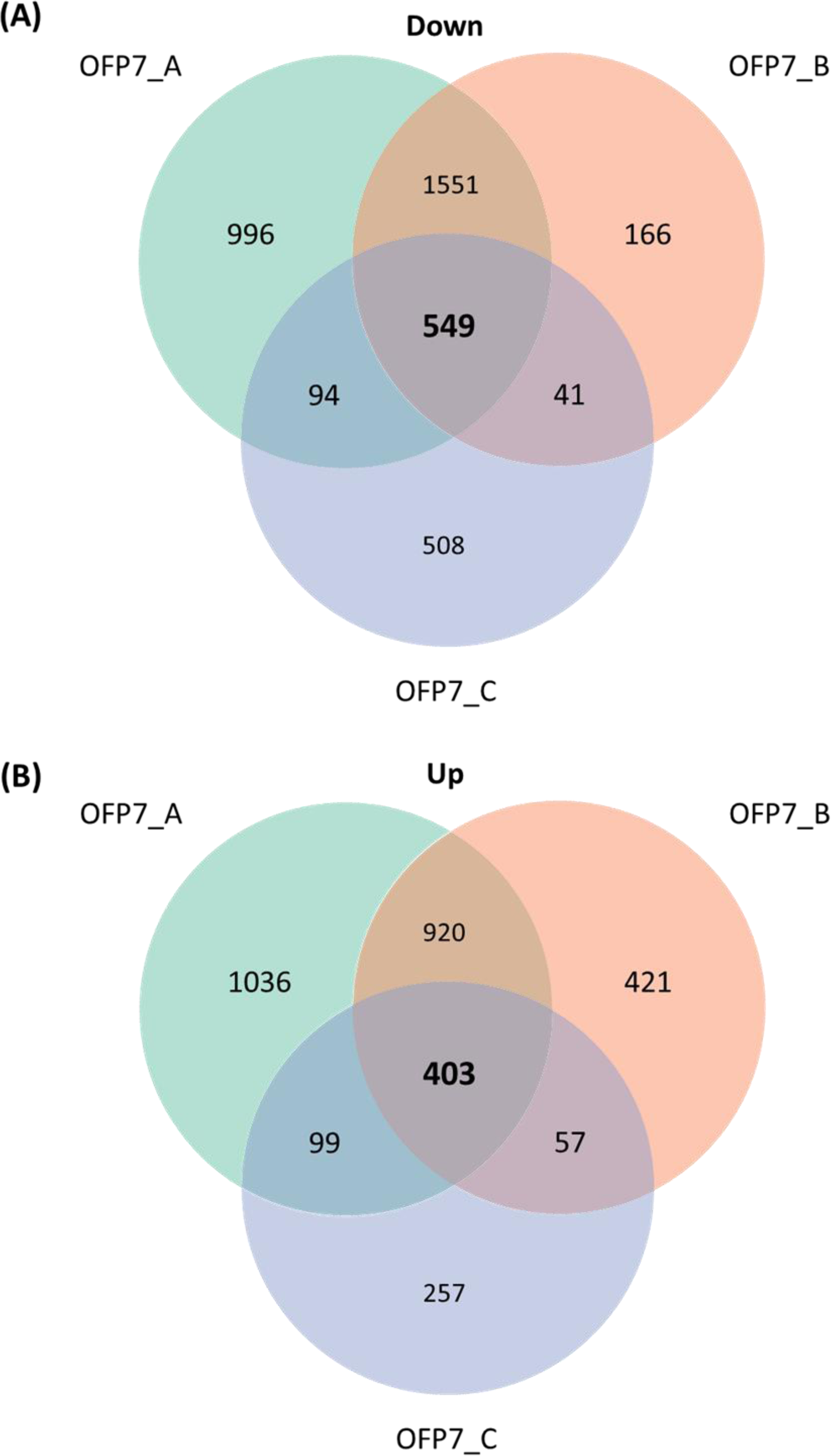
Venn diagrams showing the numbers of repressed (A) or induced (B) genes after inoculation with strains expressing dTALEs targeting PvOFP7. Differentially-expressed genes were retrieved using transcriptomic data at 2 dpi, using the EV strain as control.

### Functional analysis of differentially-expressed genes

To search for over-represented functional categories upon *PvOFP7* induction, we performed Wilcoxon statistical testing and PageMan over-representation analysis using MapMan (Fig. 6, Supplemental table 3A). Moreover, we specifically analysed important gene families (Supplemental table 4, Supplemental figure 9) including transcription factors, transcription regulators and kinases available in the iTAK database (http://itak.feilab.net) (Zheng et al. 2016), as well as *R* genes from the NLR family (Richard et al. 2017). Remarkably, *PvOFP7* was the only OFP family member to be differentially expressed, indicating that its induction did not impact other OFP genes (Supplemental figure 9).

**Fig. 6.**
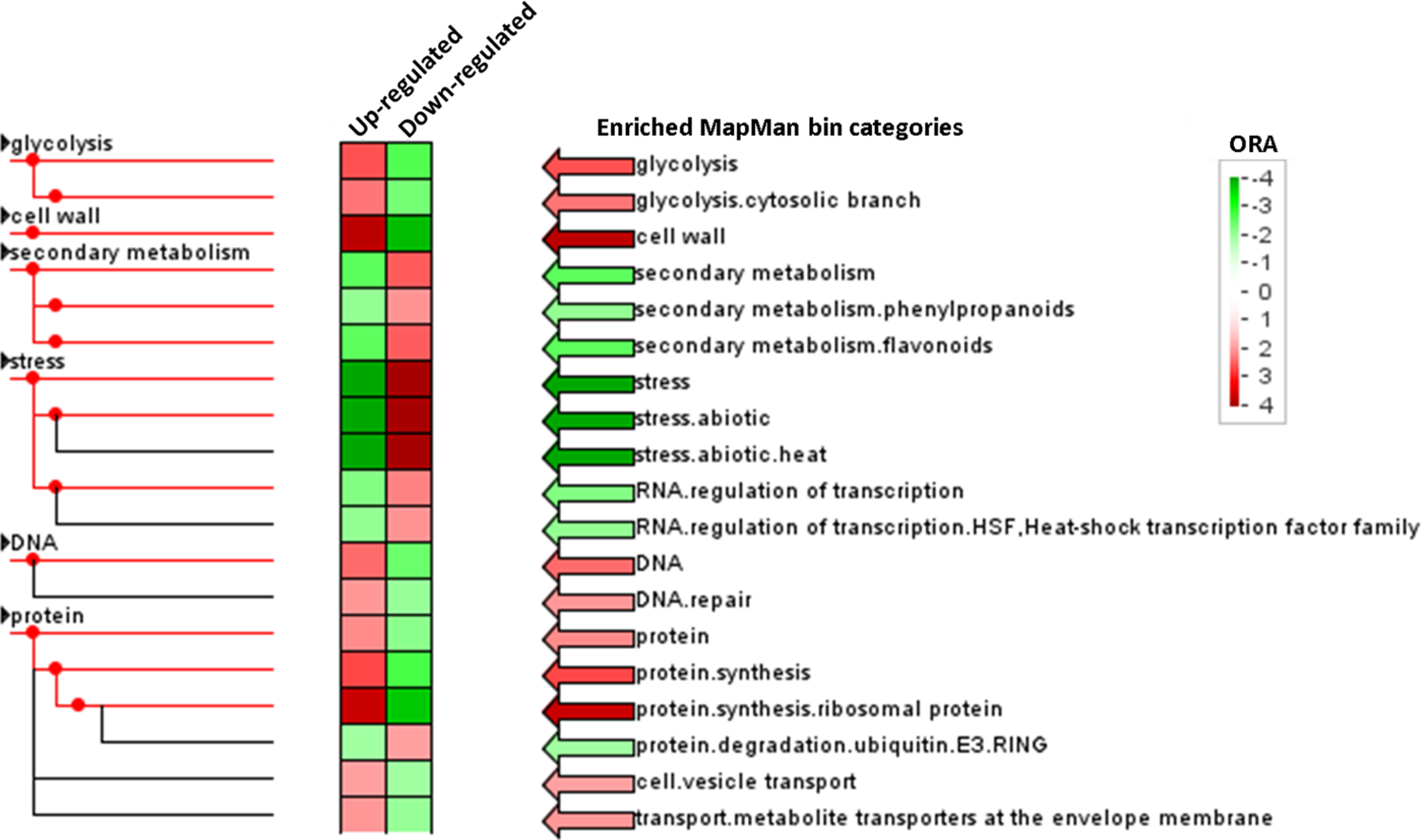
Enriched functional categories after induction of *PvOFP7.* Analysis was performed using the PageMan workflow from MapMan, on the 952 DEGs common to the three strains carrying dTALEs targeting *PvOFP7.* The colour gradient corresponds to the Over-Representation Analysis (ORA) values for up-regulated (left column) or down-regulated genes (right column). ORA values over 0 (red) correspond to over-representation while values below 0 (green) correspond to under-representation of the category.

### Down-regulation of heat shock proteins

The most significantly enriched MapMan category was related to stress (Supplemental table 3A). This category comprised a large majority of heat shock protein (HSP) genes, among which 48 out of 49 were highly repressed. In addition to HSPs, the heat shock transcription factor (HSF) family was the most impacted among all transcription factor families, with seven DEG, all down-regulated (Supplemental table 3B) indicating that repression of HSP proteins was related to *PvOFP7* induction and common bean resistance to CBB.

### Up-regulation of cell wall formation

The second most significantly enriched category encompassed genes linked to cell wall organisation, a majority of which being induced (Supplemental table 3A). This category comprised genes involved in both cell wall synthesis and cell wall modification/degradation, suggesting that cell wall remodelling occurred following *PvOFP7* induction (Supplemental table 3C). Significantly, DEGs involved in cell wall precursors and cellulose, hemicellulose and pectin synthesis, were almost all (12/14) up-regulated. In particular, we observed an up-regulation of genes involved in the production of UDP-glucose, which is a major building block of cell wall polymers (Supplemental figure 10A). Moreover, the whole pathway from Glucose-6-phosphate to UDP-xylose was induced, suggesting that biosynthesis of xylan and xyloglucan was up-regulated (Zhong *et al*., 2017). Globally, these results suggested that cell wall reinforcement occurred after *PvOFP7* induction.

### Up-regulation of primary metabolism

DEGs involved in primary functions including DNA repair, protein biosynthesis and glycolysis were mostly overexpressed, while secondary metabolism and protein degradation were down-regulated (Fig. 6). For example, ubiquitin-dependent proteolysis was down-regulated (Supplemental Figure 10B). In addition, glycolysis enzymes involved in the production of acetyl-CoA were induced, while genes involved in fermentation were repressed (Supplemental figure 10C). This suggested that the plant metabolism was oriented towards respiration via the tricarboxylic acid (TCA) cycle. Together, these observations suggested that *PvOFP7* induction impacted the plant metabolism towards primary metabolism, probably leading to production of energy and protection of important molecules such as DNA and proteins against degradation.

### Impact on plant development

Important transcription factors, regulators and kinases families such as RLKs, MYB, bHLH or bZIP were moderately impacted following *PvOFP7* induction, although with no clear pattern of induction vs. repression (Supplemental table 4). However, we observed a clear repression of genes belonging to the type B Arabidopsis Response Regulators (ARR-B) family. In Arabidopsis, *arr* mutants are impaired in cell division and exhibit altered photosynthesis and anthocyanin production (Argyros *et al*., 2008). In accordance with this, genes involved in the photosystem II and production of anthocyanins were down-regulated.

### Impact on plant defense transduction

A large majority of *R* genes from the NLR family had very low expression levels and were thus removed at the filtering step of the analysis. Following *PvOFP7* induction, only five NLR genes were differentially expressed, including a cluster of four genes of the CC-NBS-LRR subfamily that were all repressed. A gene of the TIR-NBS-LRR subfamily from the *B4* cluster was the only one that was induced (Supplemental table 4). Production of secondary metabolites such as phenylpropanoids and flavonoids was down-regulated (Supplemental figure 10D). This was quite unexpected as these metabolites are known for being key components of the plant defense (Ramaroson *et al*., 2022). Similarly, genes from the NAC family, described as key regulators of plant defenses were also repressed (Yuan *et al*., 2019). Genes from the ERF subgroup from the AP2/ERF family were down-regulated, which was consistent with the observation that they were up-regulated in a susceptible vs resistant background after CBB inoculation (Foucher *et al*., 2020). In all, *PvOFP7* did not appear to trigger a complete immune response, but rather be a key regulator of plant defense pathways downstream of primary signalling.

## DISCUSSION

In this study, we demonstrated that *PvOFP7* induction leads to strong CBB symptom reduction, in link with reduced *in planta* bacterial growth at the early stage of the colonisation. *PvOFP7* belongs to OFPs, a class of plant-specific transcriptional repressors (Wang *et al*., 2011). The first OFP to be described was OVATE, identified in tomato for its ability to cause pear-shaped fruits (Liu *et al*., 2002). OFPs were later shown to be widely distributed in the plant kingdom and to regulate multiple aspects of plant growth and development (Tsaballa *et al*., 2011; Huang *et al*., 2013; Wang *et al*., 2011; Liu *et al*., 2014; Liu *et al*., 2002). However, very little is known about the role of OFPs in plant immunity. In line with our results, two OFP genes were up-regulated in a Chinese pear genotype resistant to *Venturia nashicola* (Ding *et al*., 2020). Conversely, in cotton, GhOFP3-D13 acts as a negative regulator of plant defenses against *Verticillium dahliae* (Ma *et al*., 2020). This raises the question of how induction of an OFP gene could participate to common bean resistance to CBB.

Here, *PvOFP7* induction led to transcriptomic modifications that overlapped only partially with pathways previously described as linked to CBB resistance (Foucher *et al*., 2020). For example, we did not observe any up-regulation of the SA or down-regulation of photosynthesis and sugar metabolism, which were identified as potential markers of resistance (Foucher *et al*., 2020). Instead, we identified suppression of traits linked to susceptibility. For example, the AP2/ERF superfamily was repressed in our study while it was previously shown as being up-regulated in susceptible bean plants (Foucher *et al*., 2020). This would suggest that *PvOFP7* induction contributed to resistance by countering pathways leading to susceptibility rather than triggering a genuine defense response. In accordance with this, *PvOFP7* induction did not trigger clear up-regulation of genes involved in resistance signalling but instead led to suppression of secondary metabolites (Supplemental figure 10D), which are important defense-related phytochemicals (Piasecka *et al*., 2015).

Strikingly, *PvOFP7* induction was linked to strong repression of genes from the HSP family. This is in line with the work from Foucher *et al*. (2020), where HSP genes were strongly repressed in resistant and strongly induced in susceptible genotypes (Supplemental table 5), suggesting that suppression of HSPs is a key step towards resistance to CBB. HSPs are commonly described for their role in stress tolerance and plant immunity (Park and Seo, 2015; Sun *et al*., 2002; Ul Haq *et al*., 2019), and overexpression of HSPs is usually observed upon stress induced by pests and pathogens (Fang *et al*., 2015). Few HSPs were shown to contribute to enhanced resistance to *Xanthomonas* in pepper and rice (Kim and Hwang, 2015; Yang *et al*., 2020). In pepper, HSP70 and HSP90 were overexpressed during nonhost resistance to *X. citri* pv. *citri* (Garavaglia *et al*., 2009). However, similar to what we observed in common bean, HSPs were much more accumulated in both pepper and orange during compatible than incompatible interaction with *Xanthomonas* (Garofalo *et al*., 2009). Therefore, enhanced accumulation of HSPs could be viewed as a marker of plant susceptibility to *Xanthomonas*. Following this hypothesis, infection by the bacteria would lead to higher stress level in susceptible than resistant plant cells. In accordance with this, primary metabolism and protection against degradation were enhanced after *PvOFP7* induction, suggesting that plant cells were in a less stressed state than when confronted with the control strain.

*Xanthomonas* bacteria have developed several tactics to attack plant cells, among which targeting the integrity of the plant cell wall, which is a critical component of plant immunity (Bacete *et al*., 2018; Bartetzko *et al*., 2009). For instance, *X. oryzae* pv. *oryzae* secretes the virulence factor LipA, leading to cell wall degradation in rice (Aparna *et al*., 2009). In tomato, TALE effector AvrHah1 from *X. gardneri* triggers water-soaked lesions by indirectly up-regulating a pectate lyase (Schwartz *et al*., 2017). On the other hand, strengthening the cell wall could be critical for plant resistance to *Xanthomonas*. A perfect example of this strategy is the major resistance gene *Xa4* in rice, which encodes a cell wall-associated kinase and strengthens the cell wall by promoting cellulose synthesis and suppressing cell wall loosening (Hu *et al*., 2017). Among different functions, OFPs can be involved in secondary cell wall formation, vascular development or fruit ripening (Wang *et al*., 2016). Here, *PvOFP7* induction resulted in up-regulation of many genes involved in cell wall modification, including genes involved in the synthesis of major components of the plant cell wall such as celluloses and hemicelluloses, including xyloglucans (Eckardt, 2008; O’Neill and York, 2003; Nakano *et al*., 2015). In particular, xyloglucan cleavage, uptake and depolymerization appear as important steps for *Xanthomonas* pathogenesis (Vieira *et al*., 2021). Therefore, reinforcing the cell wall could slow down bacterial progression at the early stage of infection, resulting in reduced CBB symptoms.

Resistance to CBB is known for being complex and accumulation of multiple resistance QTLs is needed to acquire strong resistance to this disease (Chen *et al*., 2021). This is the case for the resistant cv. BAT93, that cumulated four resistance QTLs to CBB (Nodari *et al*., 1993) and from which we unveiled *PvOFP7* as a candidate for resistance (Foucher *et al*., 2020). *PvOFP7* is located in a resistance QTL on chromosome 9, which explains 13% of the phenotypic variation (Nodari *et al*., 1993). Therefore, *PvOFP7* is only a partial component of common bean defense to CBB, and other genes might contribute to CBB resistance. Significantly, none of the other 13 candidate genes tested in this study were differentially expressed following *PvOFP7* induction, this included *PvAP2-ERF71* and *PvExpansinA17*, whose induction also led to symptom reduction. This would suggest that these genes operate in other pathways independent from *PvOFP7*. However, joint induction of *PvOFP7* and *PvExpansinA17* led to negative regulation of *PvAP2-ERF71*, reflecting complex relationships between these three genes.

Partial resistance was achieved by specifically inducing *PvOFP7* in susceptible genotypes. However, *PvOFP7* transcription levels after dTALEs induction in JaloEEP558 were somewhat extreme compared to what was previously observed in BAT93 inoculated with the 6546R strain (Foucher *et al*., 2020). Moreover, PvOFP7-mediated resistance was dose-dependent. Thus, we believe that the symptom reduction of more than 60% observed here is an overestimation of what may happen during natural infection, where one would expect more subtle changes in *PvOFP7* transcriptional regulation. Indeed, *PvOFP7* promoter region was not 100% identical between JaloEEP558 and BAT93 (pairwise percentage identity of 98.7% over 1,000 bp) (Supplemental figure 11), suggesting that differences in *PvOFP7* regulation may exist that could explain resistance *vs* susceptibility. In this regard, it would be interesting to investigate whether genetic and/or epigenetic polymorphism could differentiate resistant from susceptible common bean accessions.

Our study highlighted *PvOFP7* as a major contributor to CBB resistance. This is, to our knowledge, the first functional description of a gene involved in CBB resistance. Remarkably, we pointed out a correlation between *PvOFP7* induction and symptom reduction, which makes sense with resistance to CBB being quantitative, but also shows the remarkable ability of the dTALE approach to unveil phenotypes that would be impossible to detect by knock-out mutagenesis. Although more in-depth validation is needed for *PvAP2-ERF71* and *PvExpansinA17*, these findings provide valuable insights into three genes that may be useful for CBB bean resistance breeding.

## ACKNOWLEDGEMENT

We deeply thank Dr Jérôme Verdier (IRHS, France) for his help in RNA-Seq analyses. We thank Pierre Santagostini and Angelina El Ghaziri (IRHS, France) for the useful discussions about statistics. Bacterial strains were handled at the CIRM-CFBP (CIRM-CFBP, 2021). This study was supported by the French National Research Agency project SUCSEED (ANR-20-PCPA-0009). CG was funded for her PhD by the French Pays de Loire Region, the French National Research Institute for Agriculture and Environment, Plant Health and Environment division (INRAE-SPE) and L’Institut Agro Rennes-Angers.

## AUTHOR CONTRIBUTIONS

**Charlotte GAUDIN**: Conceptualization, Formal analysis, Investigation, Methodology, Supervision, Visualization, Writing - Original Draft. **Anne PREVEAUX**: Formal analysis, Investigation, Methodology, Visualization. **Nathan AUBINEAU**: Investigation. **Damien LE GOFF**: Investigation. **Marie-Agnès JACQUES**: Conceptualization, Supervision, Writing - Review & Editing. **Nicolas W.G. CHEN**: Conceptualization, Funding acquisition, Project administration, Writing - Original Draft, Writing - Review & Editing.

## DATA AVAILABILITY

Raw RNA-seq data were deposited at the NCBI SRA database under BioProject ID <u>PRJNA1000747</u>. Bacterial strains are available upon request to Nicolas W.G. Chen.

