## Supplemental figure 1 for "Induction of common bean *OVATE Family Protein 7* (*PvOFP7*) promotes resistance to common bacterial blight"

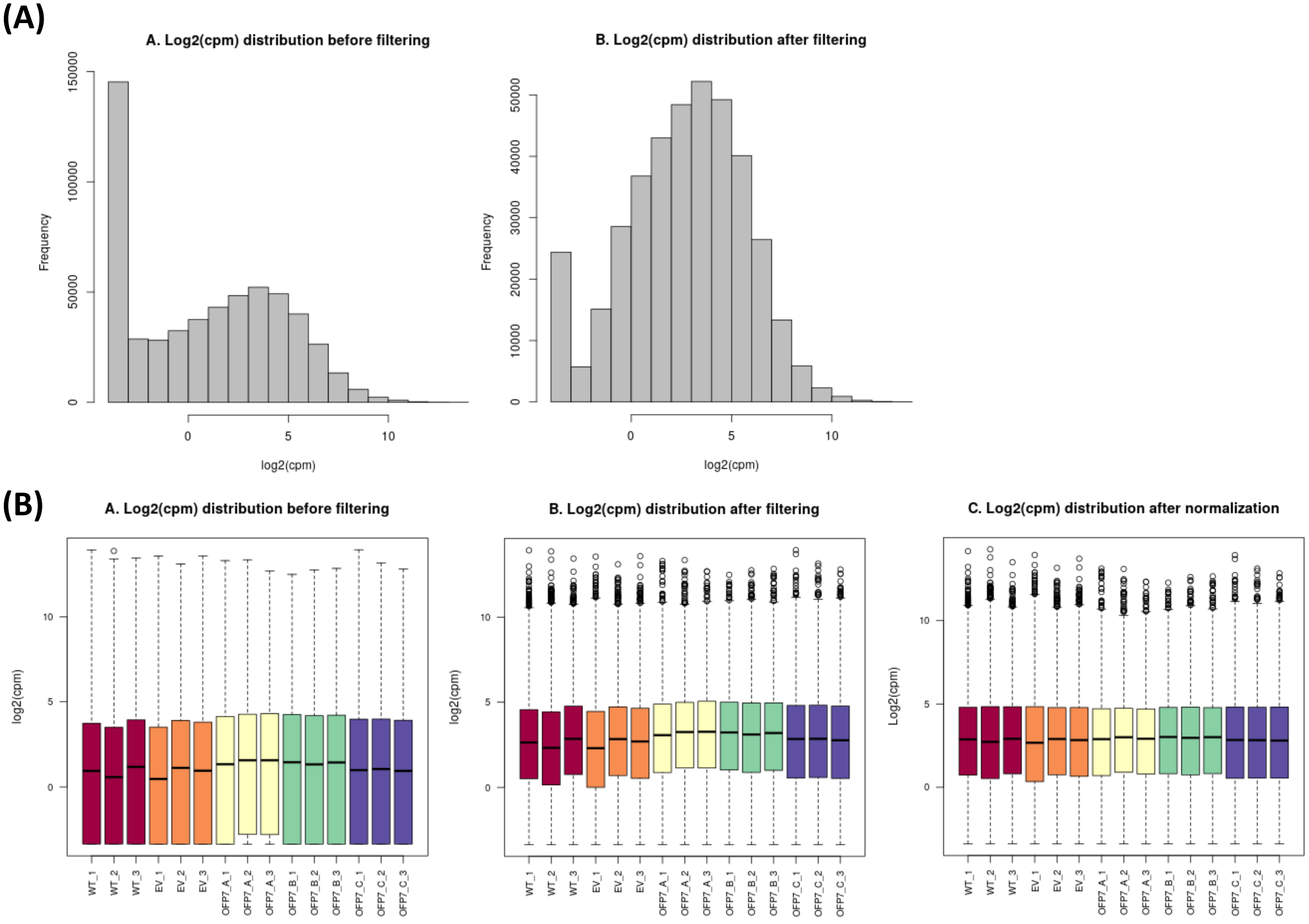


**Supplemental figure 1.** Statistics showing the effect of filtering and normalization for RNA-Seq data. **A)** Frequency distribution of $Log_2$(CPM) before (left) and after (right) filtering. **B)** $Log_2$(CPM) values before (left), after (middle) filtering and after normalization (right).
