## Supplemental figure 2 for "Induction of common bean *OVATE Family Protein 7* (*PvOFP7*) promotes resistance to common bacterial blight"

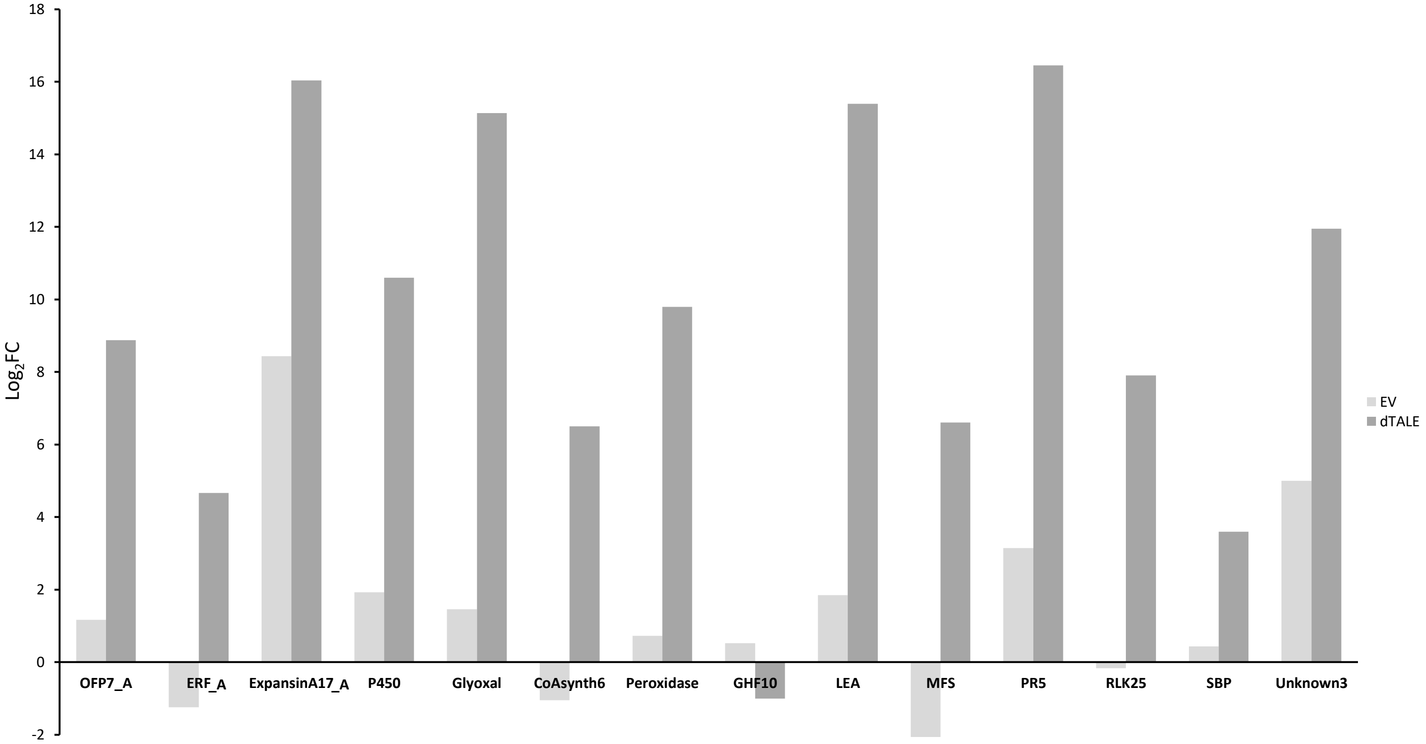


**Supplemental figure 2.** *In planta* induction of 14 candidate genes using dTALEs. Trifoliate leaves were vacuum-infiltrated at $1.10^8$ cfu.ml$^{-1}$ and harvested 48 hours post-inoculation to run qRT-PCRs. $Log_2$(FC) was calculated using the $2^{(-\Delta\Delta CT)}$ method on the plants inoculated by the EV strain as controls and the *IDE* gene as internal control. Means were calculated on data from three individuals per treatment. Equivalences between dTALE names and gene IDs are found in Supplemental table 1.
