## Supplemental figure 3 for "Induction of common bean *OVATE Family Protein 7* (*PvOFP7*) promotes resistance to common bacterial blight"

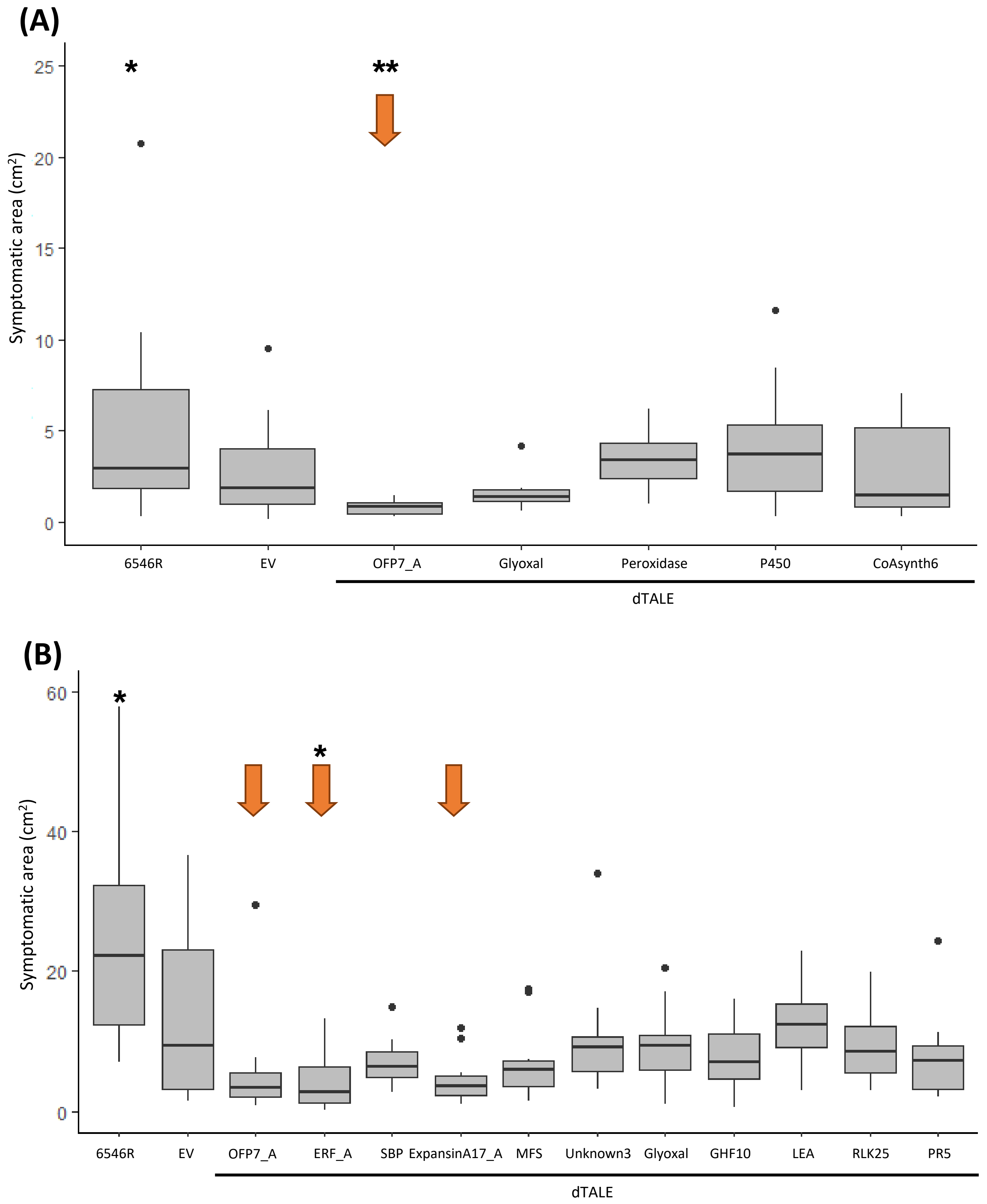


**Supplemental figure 3.** Screening of 14 candidate genes for CBB symptom reduction. First leaves of JaloEEP558 were rub-inoculated with bacterial suspensions calibrated at $1\times10^{8}$ cfu.mL$^{-1}$. A and B correspond to two independent experiments. Average symptomatic areas were assessed at 9 (A) and 10 dpi (B). Ten plants were used per treatment. Kruskal Wallis test indicated significant effect ($p<0.05$) of treatments. The number of stars refers to significantly different values compared to the EV after Dunn's test (* < 0.05, ** < 0.01). Arrows show the three most interesting candidate genes used in Figure 1. Equivalences between dTALEs name and genes ID are found in the supplemental table 1.
