## Supplemental figure 4 for "Induction of common bean *OVATE Family Protein 7* (*PvOFP7*) promotes resistance to common bacterial blight"

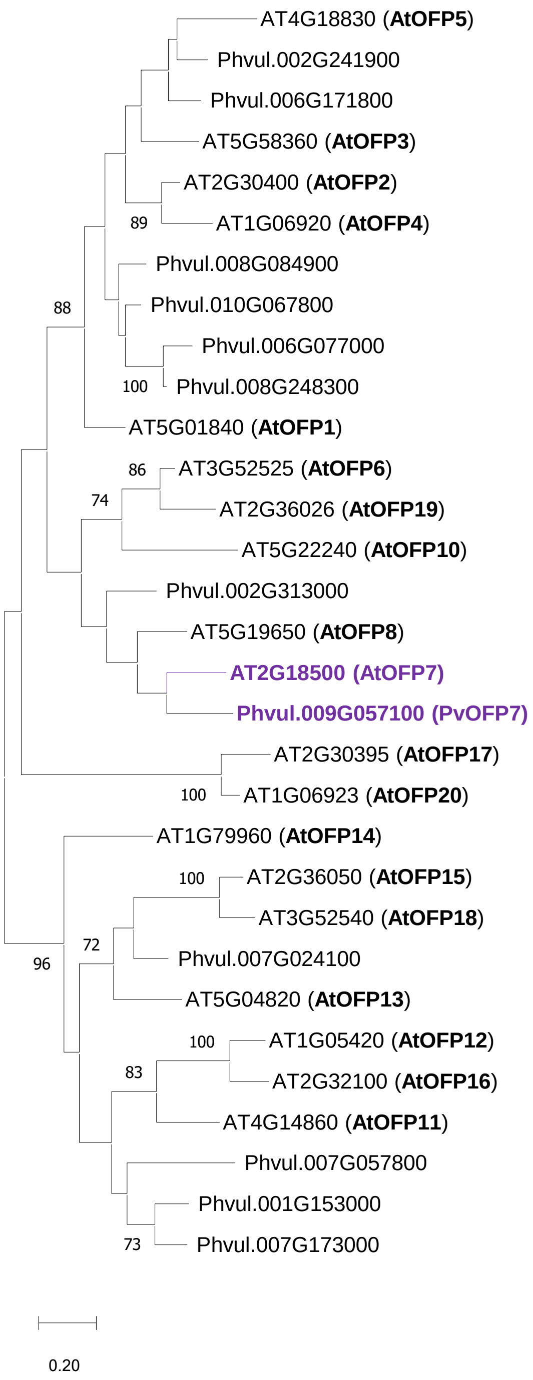


**Supplemental figure 4.** Phylogeny of OFPs from *Arabidopsis thaliana* and *Phaseolus vulgaris*. *PvOFP7* and *AtOFP7* are shown in purple. This maximum Likelihood tree was done using amino acid sequences of OVATE domains of all *A. thaliana* and *P. vulgaris* genes. Number at nodes represent bootstrap values for 100 iterations, with a threshold of 70, and the horizontal scale bar represents the number of nucleotide substitutions per site.
