## Supplemental figure 5 for "Induction of common bean *OVATE Family Protein 7* (*PvOFP7*) promotes resistance to common bacterial blight"

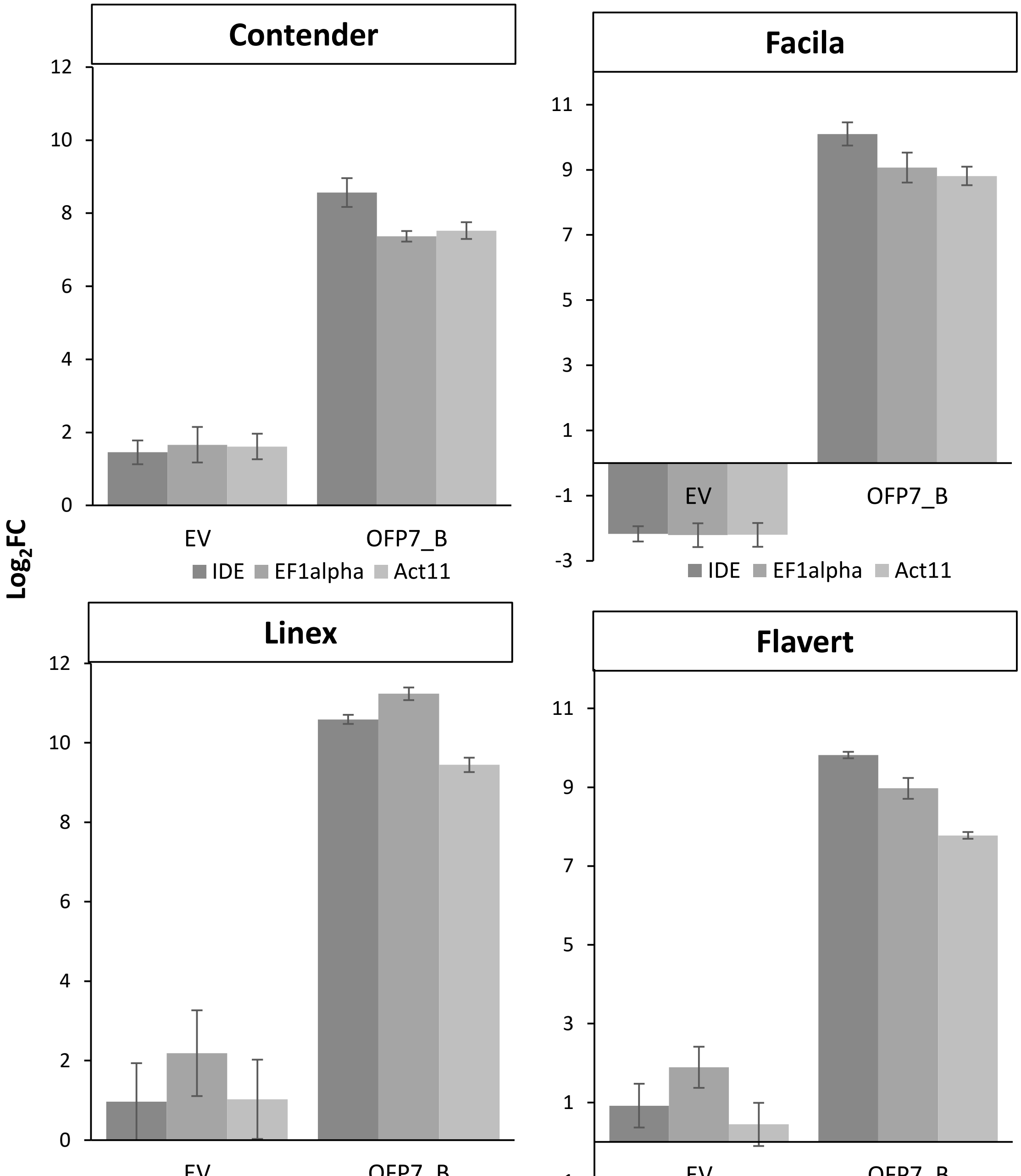


**Supplemental figure 5.** *PvOFP7* expression levels measured by qRT-PCR at 2 dpi in four additional common bean accessions. Trifoliate leaves were vacuum-infiltrated at $1x10^8$ cfu.ml$^{-1}$. $Log_2$(FC) were calculated using the $2^{(-\Delta\Delta CT)}$ method control. These results come from two independent experiments, each comprising three samples per treatment (n=6).
