## Supplemental figure 6 for "Induction of common bean *OVATE Family Protein 7* (*PvOFP7*) promotes resistance to common bacterial blight"

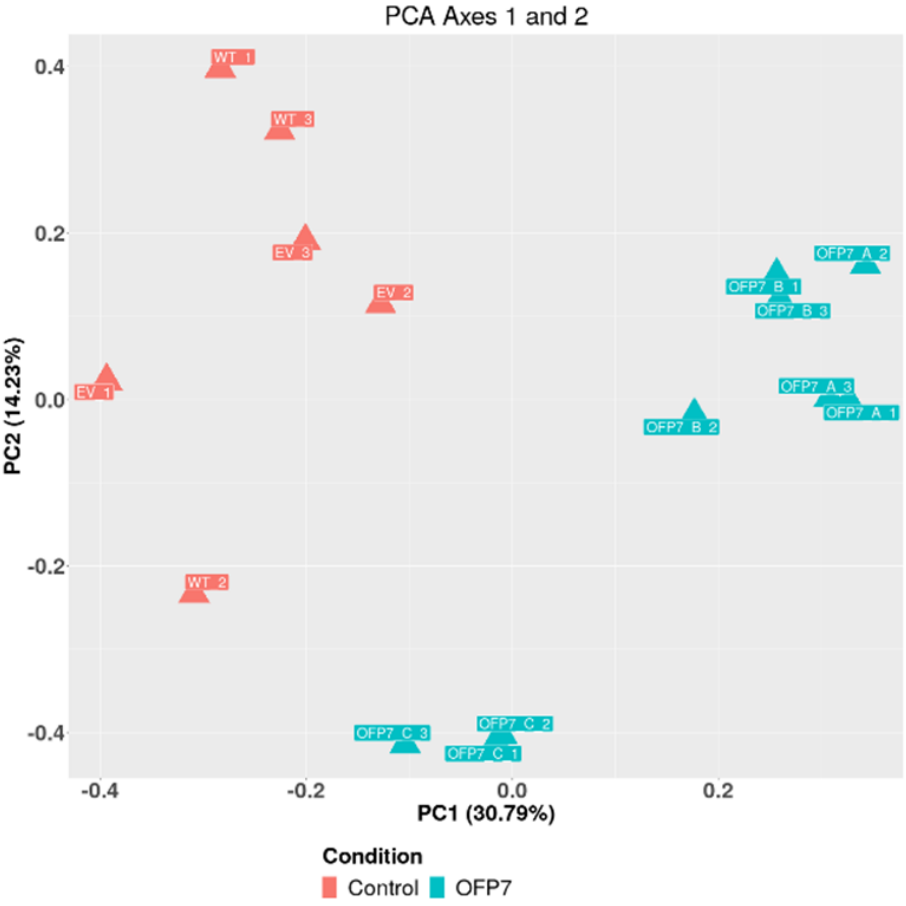


**Supplemental figure 5.** Principal Component Analysis (PCA) plot showing the variance of mapped reads between common bean leaves inoculated with the control strains and the strains carrying dTALEs. This analysis was made with AskoR with the first two principal axes explaining 30.79% and 14.23% of the variance. Orange triangles represent the controls inoculated with strains 6546R or EV. Blue triangles represent the conditions corresponding to strains expressing dTALEs OFP7_A, OFP7_B and OFP7_C, targeting different regions of the PvOFP7 promoter.
