## Supplemental figure 7 for "Induction of common bean *OVATE Family Protein 7* (*PvOFP7*) promotes resistance to common bacterial blight"

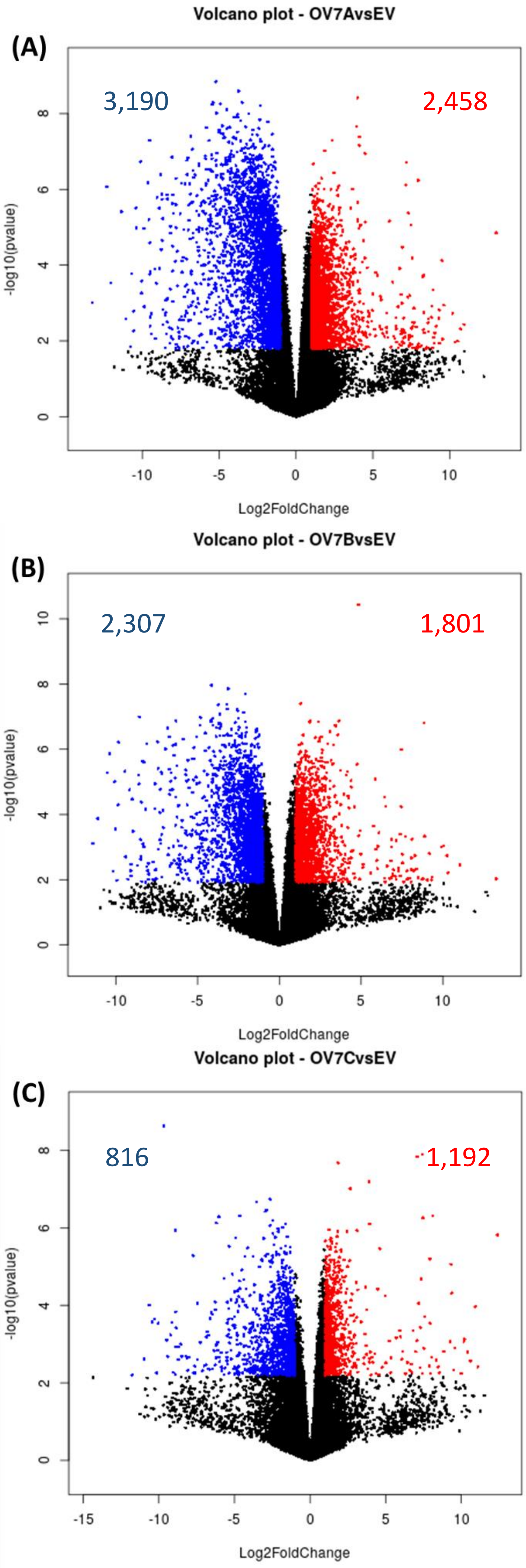


**Supplemental figure 7.** Global impact of *PvOFP7* induction on the common bean transcriptome. Volcano plots represent the distribution of DEGs in common bean plants inoculated with strains caring dTALEs OFP7_A **(A)**, OFP7_B **(B)** or OFP7_C **(C)** using the EV strain as control. Blue dots indicate down-regulated genes while red dots correspond to up- regulated genes.
