## Supplemental figure 8 for "Induction of common bean *OVATE Family Protein 7* (*PvOFP7*) promotes resistance to common bacterial blight"

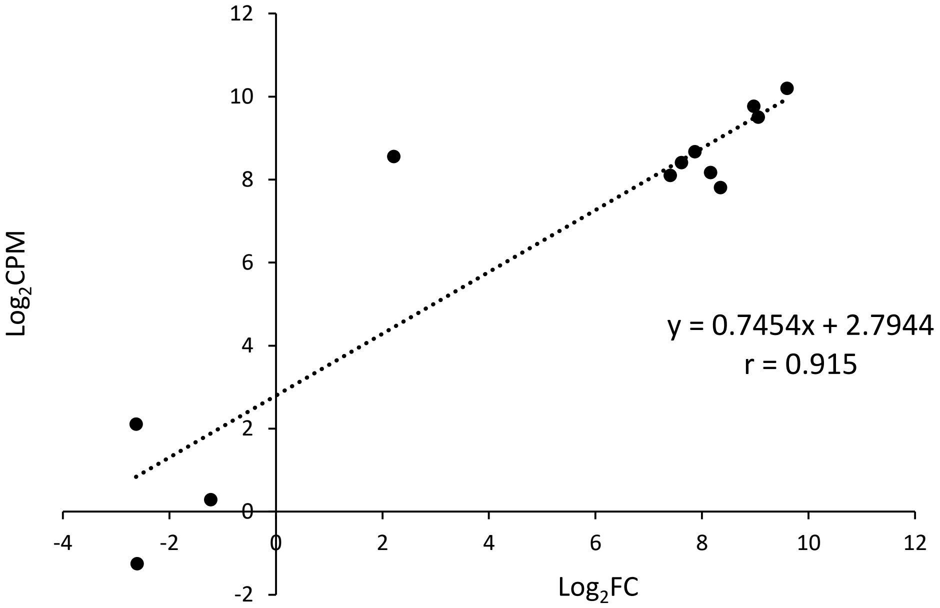


**Supplemental figure 8.** Linear regression between qRT-PCR and RNA-Seq results for *in planta PvOFP7* expression levels. Each dot corresponds to one sample for which *PvOFP7* expression levels were measured using both qRT-PCR and RNA-Seq, in plants inoculated with EV and dTALEs-complemented strains. Correlation coefficient of +0.91 was significant ($p<0.001$) for $n=12$ observations.
