## Supplemental figure 9 for "Induction of common bean *OVATE Family Protein 7* (*PvOFP7*) promotes resistance to common bacterial blight"

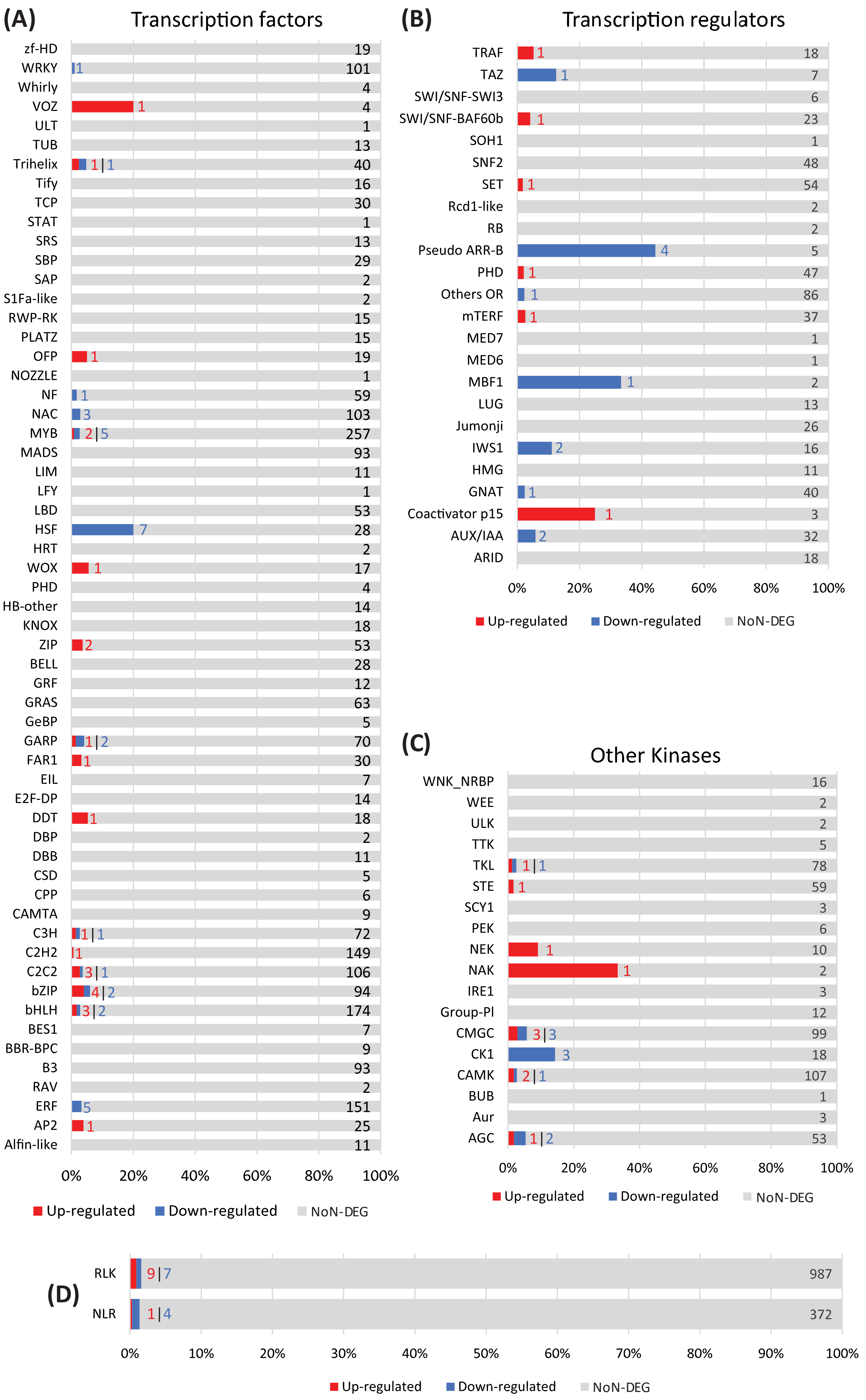


**Supplemental figure 9.** Proportion of differentially expressed transcription factors **(A)**, transcription regulators **(B)**, other kinases **(C)**, RLK and NLR **(D)**. Up-regulated genes are indicated in red and down-regulated genes in blue. Genes that are either not represented in our data or with an FDR > 0.05 are represented in grey. The number of genes in each category is indicated on the bar chart.
