## Supplemental figure 10 for "Induction of common bean *OVATE Family Protein 7* (*PvOFP7*) promotes resistance to common bacterial blight"

**(A)**

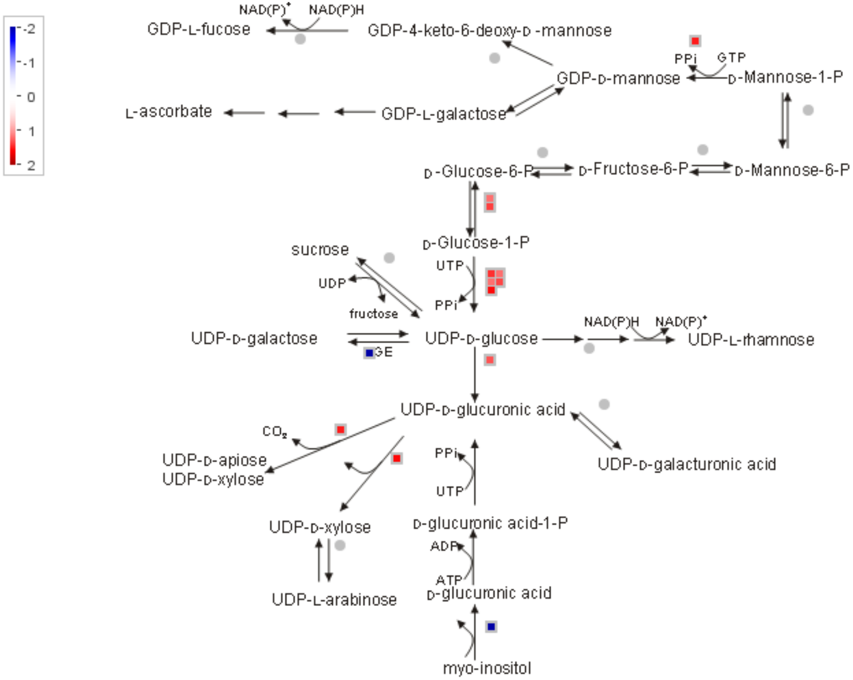


**Supplemental figure 10.** MapMan representations of DEGs within pathways related to cell wall precursors **(A)**, the proteasome **(B)**, glycolysis **(C)** or secondary metabolites **(D)**. The colour gradient indicates $Log_2$(FC) level of expression.

**(B)**

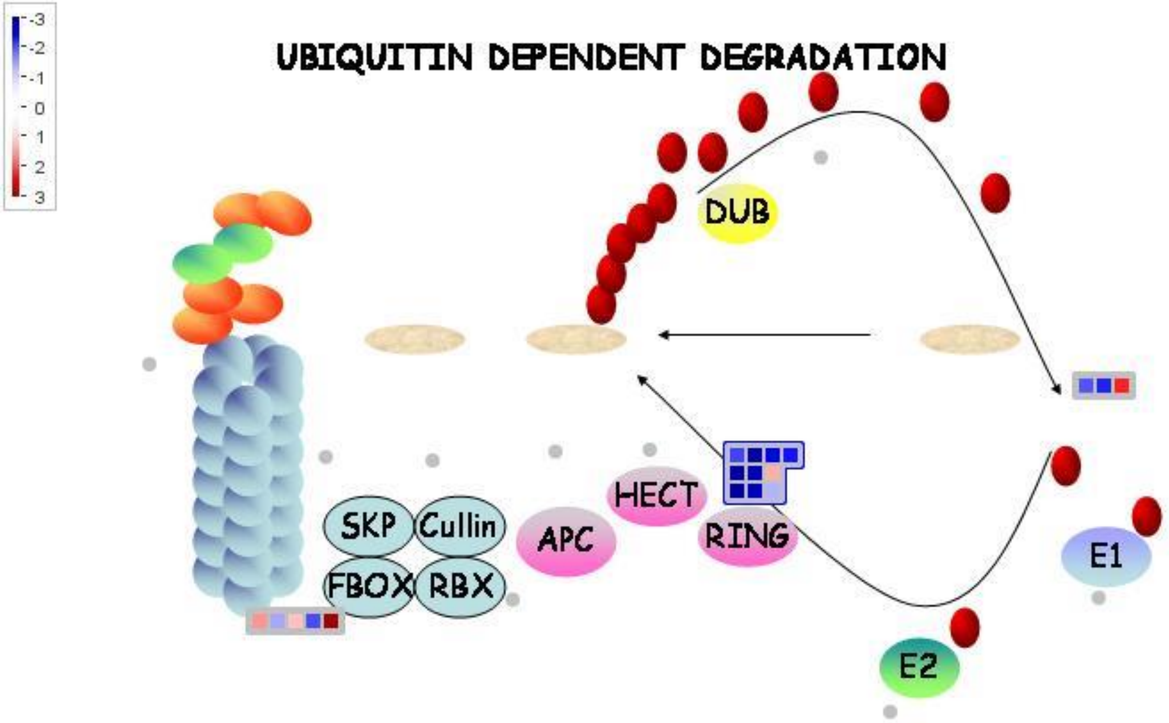


**Supplemental figure 10.** MapMan representations of DEGs within pathways related to cell wall precursors **(A)**, the proteasome **(B)**, glycolysis **(C)** or secondary metabolites **(D)**. The colour gradient indicates $Log_2$(FC) level of expression.

**(C)**

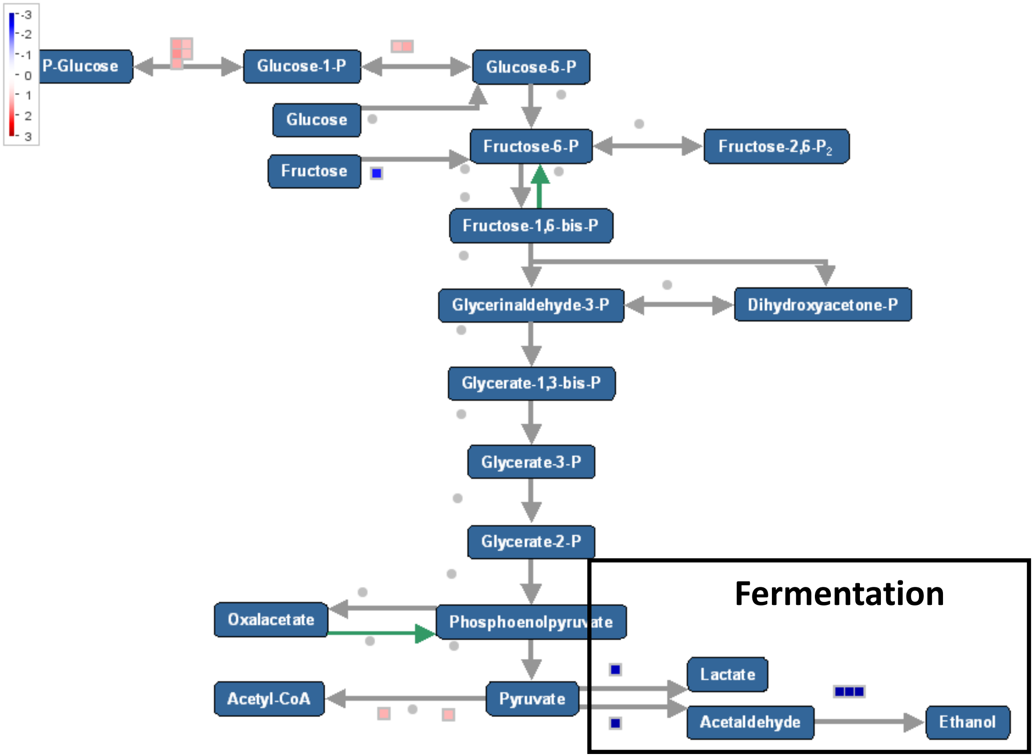


**Supplemental figure 10.** MapMan representations of DEGs within pathways related to cell wall precursors **(A)**, the proteasome **(B)**, glycolysis **(C)** or secondary metabolites **(D)**. The colour gradient indicates $Log_2$(FC) level of expression.

(D)

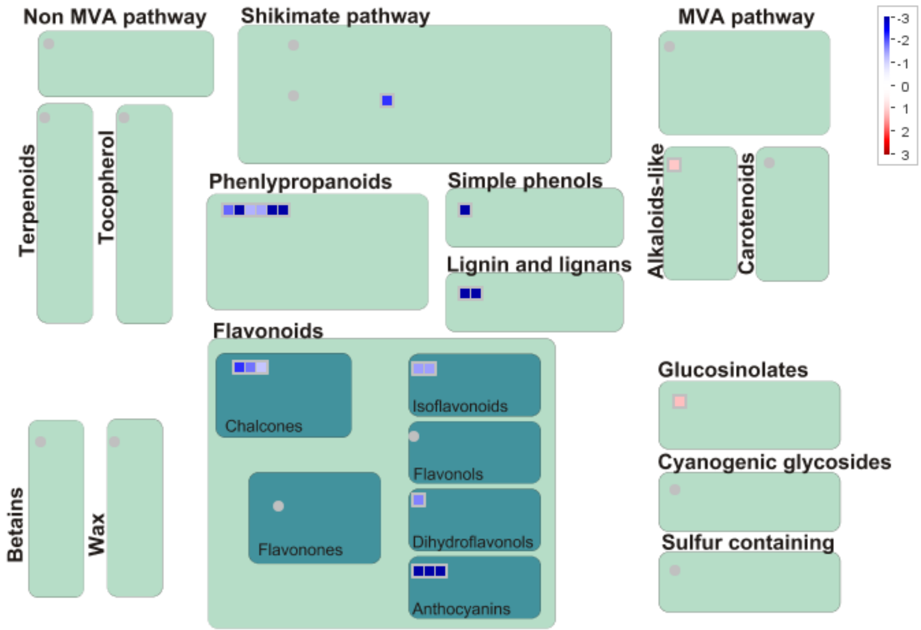


**Supplemental figure 10.** MapMan representations of DEGs within pathways related to cell wall precursors **(A)**, the proteasome **(B)**, glycolysis **(C)** or secondary metabolites **(D)**. The colour gradient indicates $Log_2$(FC) level of expression.
