## Supplemental figure 11 for "Induction of common bean *OVATE Family Protein 7* (*PvOFP7*) promotes resistance to common bacterial blight"

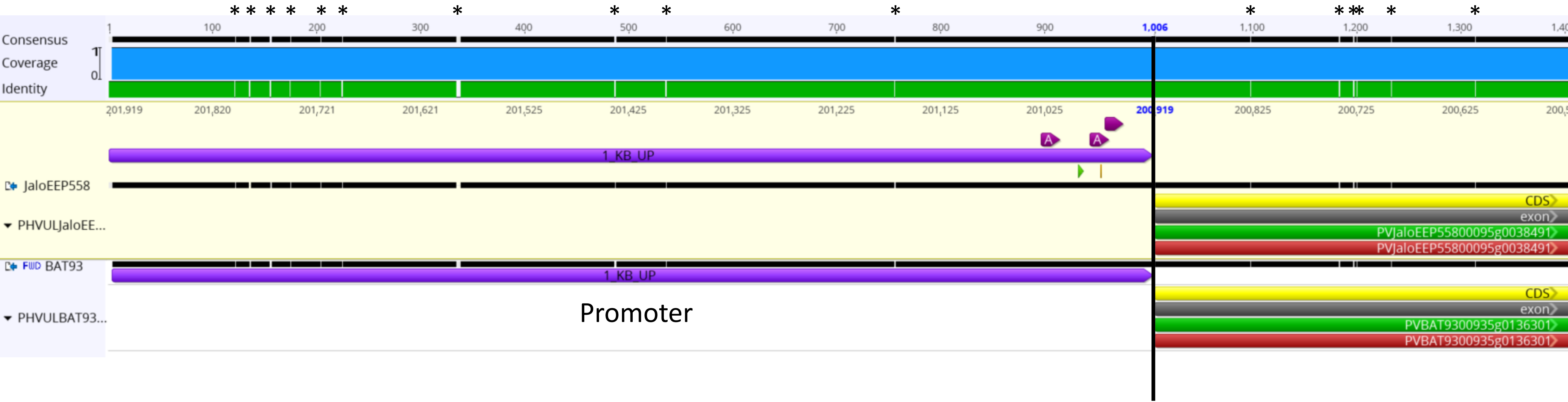


**Supplemental figure 11.** Comparison of *PvOFP7* promoter regions of BAT93 and JaloEEP558 visualized using Geneious. Asterisks indicate polymorphic sites.
